# Characterization of Inter-Alpha-Trypsin Inhibitor Heavy Chain 5 (ITIH5) Expression, a Prostate Basal Cell Marker

**DOI:** 10.64898/2026.11.13.766426

**Authors:** Gopal Kumar, Darshan Shimoga Chandrashekar, Sofia Cañete-Portillo, Farrukh Afaq, Shannon Carskadon, Upender Manne, George J. Netto, Edgar Dahl, Nallasivam Palanisamy, Sooryanarayana Varambally

**Affiliations:** Department of Pathology, University of Alabama at Birmingham, Birmingham, AL, USA; Vattikuti Urology Institute, Department of Urology, Henry Ford Health System, Detroit, Michigan; Department of Pathology and Laboratory Medicine Perelman School of Medicine University of Pennsylvania, Philadelphia, Pennsylvania; Institute of Pathology, Medical Faculty, RWTH Aachen University, 52074, Aachen, Germany; Center for Integrated Oncology Aachen Bonn Cologne Düsseldorf (CIO ABCD), 52074, Aachen, Germany; O’Neal Cancer Center, University of Alabama at Birmingham, Birmingham, AL, USA

## Abstract

The prostate epithelium contains secretory luminal epithelial cells, basal epithelial cells and neuroendocrine cells. Luminal cells are secretory cells that produce prostatic secretions and basal cells are involved in regeneration and maintenance of epithelial tissue. The absence of a basal cell layer is a defining morphological feature of prostate adenocarcinoma. Immunohistochemical detection of basal cells using standard markers such as p63 and high molecular weight keratin (34βE12) is routinely employed to differentiate invasive carcinoma from benign architecture mimics. Additional protein like TRIM29 and ETS2 are also known to stain prostatic basal cells. While recent advances in sequencing and proteomic technologies helped identify many luminal epithelial markers for prostate cancers, basal cells are less investigated. In the current study, we identified Inter-Alpha-Trypsin Inhibitor Heavy Chain 5 (ITIH5) as a gene expressed exclusively in the basal cells of prostate epithelium. ITIH5 is a hyaluronan-stabilizing extracellular matrix protein that has been shown to be downregulated by promoter DNA hypermethylation in various cancers. Gene expression analysis using published dataset suggested ITIH5 is expressed in prostate basal cells compared to luminal cells. To characterize the potential diagnostic utility of ITIH5 in prostatic tissues, we evaluated its immunohistochemical expression profile across benign and malignant prostatic glands. Our immunohistochemistry analysis confirmed that ITIH5 protein is mainly in the cytoplasmic compartment of basal cells lining benign prostatic acini while prostate luminal cells showed no signal for ITIH5. Importantly, the cytoplasmic staining of ITIH5 reflected the nuclear staining of p63 in terms of cell they are staining, a well-established prostate basal cell marker that is a nuclear protein. Our results suggest ITIH5 serves as a novel basal cell marker for prostate tissue differential diagnosis on its own or in combination with other established markers. Biological role and significance of ITIH5 expression in prostate basal cells need to be elucidated.

## Introduction

The normal prostate epithelium is composed of three distinct cell types: secretory luminal cells, basal cells, and neuroendocrine cells (1). While secretory luminal cells are functional, androgen-dependent epithelial cells that line the glandular ducts and acini of the prostate, basal cells form a boundary between stroma and the luminal cells (2). Prostate neuroendocrine cells make a small population of cells that are involved in secreting growth factors and hormones(3). In diagnostic surgical pathology, the evaluation of the basal cell compartment is fundamentally important because the malignant transformation of prostate acini is characteristically associated with the absence of the basal cell layer (4). Usual type (acinar) prostatic adenocarcinoma, which comprises majority of prostate cancers, originates from the luminal secretory cells of the prostatic acini and ducts (5). Prostate cancers may also contain neuroendocrine cells(6). The absence of basal cells is one of the hallmarks of prostatic adenocarcinoma (7). While the process of loss of basal cells in malignant prostate cancer is still not very well understood, current standard diagnostic algorithms rely heavily on established basal cell markers, like high molecular weight cytokeratins (recognized by the 34βE12 antibody) and the nuclear protein p63(8). While 34βE12 targets cytoplasmic and membranous antigens, p63 is a nuclear transcription factor(9). In clinical practice, these markers are frequently utilized together in a basal cell cocktail to maximize detection sensitivity, as they target distinct subcellular compartments. Consequently, demonstrating the loss of basal cells via immunohistochemistry is a critical step in distinguishing well-differentiated prostatic adenocarcinoma from benign architectural mimics(8–10). Despite the increased utility of these immunohistochemical markers, diagnostic challenges still persist, for example, certain benign lesions such as adenosis, can exhibit inconspicuous or focally absent basal cell layer(9). In addition, certain technical factors like tissue fixation and antigen retrieval methods may lead to false-negative staining or variable expression of current markers (9). Given these limitations, the identification and characterization of novel, highly specific basal cell markers are clinically important in the differential diagnosis of atypical prostatic proliferations. Our current study demonstrates that ITIH5, a member of the inter-a-trypsin inhibitor heavy chain family, originally identified as downregulated in breast cancer (11,12) and a tumor and metastasis suppressor involved in extracellular matrix stability, the loss of which may facilitate stromal invasion and metastasis (13)(14,15)(16–18), is robustly expressed in prostatic basal cells and absent from invasive prostate cancer glands. While loss of ITIH5 expression due to promoter DNA hypermethylation is common in various cancers including breast, bladder, lung, colorectal, and cervical cancer(19) (20–22) (11,26) (23), its expression has not been investigated in prostate cancer.

## Materials and methods

### Gene expression analysis

Prostatic basal cell gene expression analysis was performed using the publicly available data set GSE67070(24). Briefly, raw read count data for all six samples [Prostatic Basal cells :GSM1638209, GSM1638210, GSM1638211; Prostatic Luminal cells: GSM1638212, GSM1638213, GSM1638214] from this dataset were downloaded from the NCBI Gene Expression Omnibus (GEO) repository. Differential gene expression analysis was performed in R (version 4.5.2) using the DESeq2 package [PMID: 25516281]. Volcano plots were generated using the EnhancedVolcano R package to visualize differentially expressed genes, and jitter-box plots were created using the ggplot2 package to display gene expression distributions across sample groups [https://github.com/kevinblighe/EnhancedVolcano; https://ggplot2.tidyverse.org]. All statistical analyses and data visualizations were performed within the R 4.5.2 environment.

### Immunohistochemistry

P63 mouse monoclonal antibody (Biocare Medical, SKU: 163) and ITIH5 rabbit polyclonal antibody generated by Pineda Antibody Service, Berlin, raised against a C-terminal ITIH5 peptide and characterized previously (15) were used for the immunostaining of prostate tissues. Immunohistochemistry was performed on PCa cancer tissue specimens (Institutional review board approved). Briefly, tissue sections were deparaffinized, rehydrated, and antigen retrieval was performed at 100°C for 5 minutes in citrate buffer (pH 6.0), followed by blocking with 5% Normal Goat Serum, and incubation with primary antibodies (ITIH5, Anti-rabbit, 1:200 dilution, p63, Anti-mouse, 1:100, Biocare Medical, SKU: 163) for 1 hour at room temperature and with goat anti-Rabbit (abcam: ab6721, 1/1000 for ITIH5 and Goat anti-Mouse (abcam: ab6789) 1/200 for p63) for 1 hour at room temperature. After washing the slide in TBST and TBS buffers, diaminobenzidine staining was performed (Vector Laboratories - ImmPACT DAB Peroxidase (HRP) Substrate Kit/SK4105 (Cat# SK-4105, Vector laboratories) was used to assess immunoreactivity, and counter-stained with Hematoxylin 1 (Fisher Scientific 22-050-112). Images are acquired using Aperio CS2 Scanner (Leica Biosystems).

### Immunofluorescence staining

Immunofluorescence staining was performed on prostate tissue with ITIH5 rabbit polyclonal antibody (13). Briefly, tissue sections were deparaffinized, rehydrated, and antigen retrieval was performed at 100°C for 5 minutes in citrate buffer (pH 6.0), followed by blocking with 5% Normal horse Serum (Vector Biolab, S-2012-50, and incubation with primary antibodies for 2 hours at room temperature (ITIH5, Anti-rabbit, 1:400 dilution). After washing with TBST, the slides were incubated with Donkey anti-Rabbit IgG (H+L) Secondary Antibody conjugated to Alexa Fluor™ Plus 488 (ThermorFIsher Scientific, A32790), at 1:1000 dilution for 1 hours at room temperature. After washing with TBST and TBS the ProLong™ Gold Antifade Mountant with DNA Stain DAPI was added and coverslip was mounted and the sides were sealed. The stained slides were imaged using Leica Microsystem, Model Stellaris V Confocal Micrscope and used z-stacks.

## Results

In order to evaluate basal cell markers, we used the publicly available and previously published gene expression dataset to perform differential gene expression analysis (GSE67070, Ref, PMID: 26924072). This analysis identified the known, published prostate basal cell markers like p63, ETS2, TRIM29. Interestingly, our analysis also showed thatInter-Alpha-Trypsin Inhibitor Heavy Chain 5 (ITIH5) is enriched in prostate basal cells compared to luminal cells (**Fig. 1A, B**).

**Figure 1:**
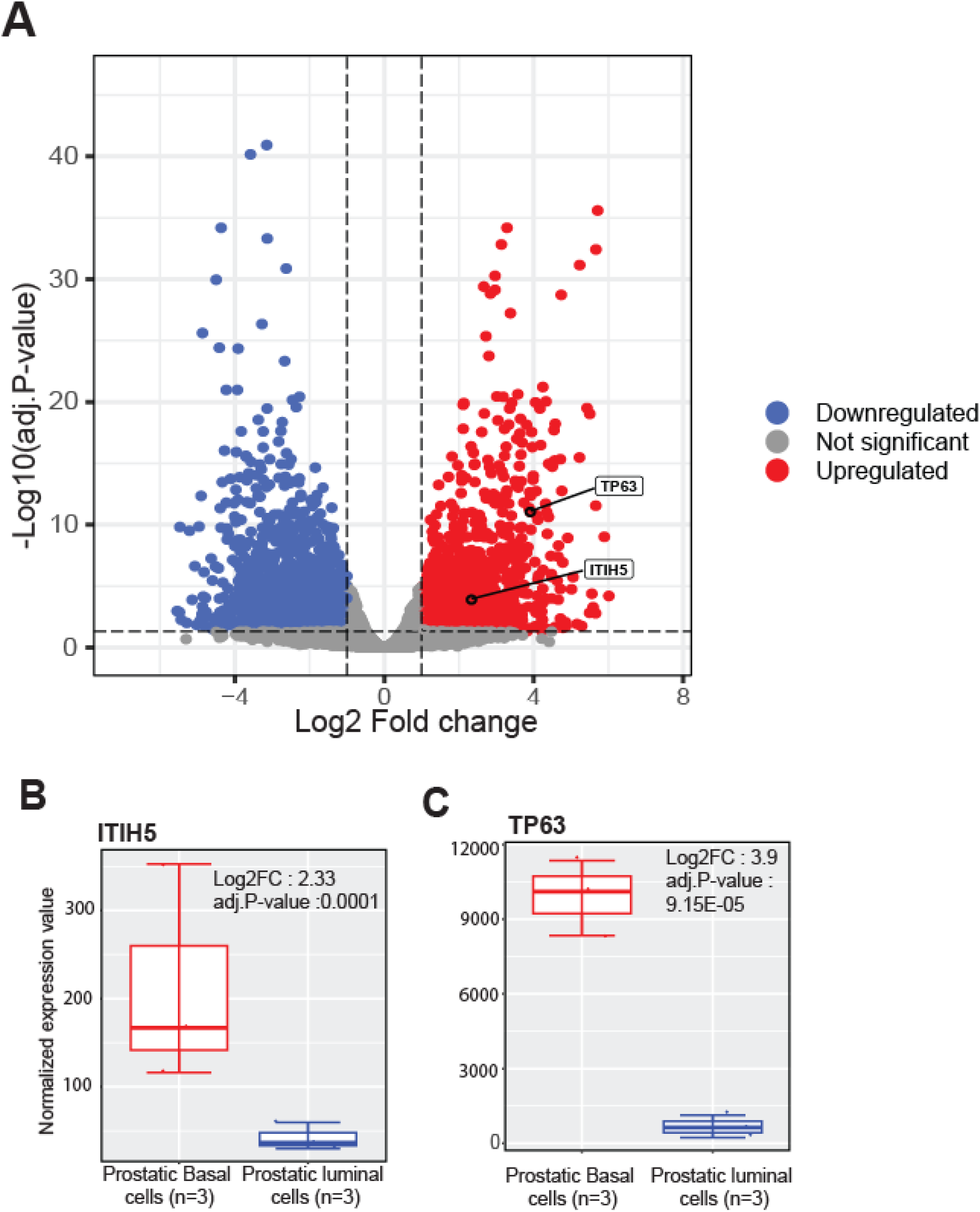
Differential Gene Expression Analysis Between Prostatic Basal and Luminal Cells. (A) Volcano plot showing differentially expressed genes between prostatic basal cells (n = 3) and prostatic luminal cells (n = 3). Differentially expressed genes were identified using thresholds of |log□ fold change| ≥ 1 and adjusted P-value < 0.05. Upregulated genes are shown in red, downregulated genes in blue, and non-significant genes in gray. (B–D) Jitter-box plots showing the expression levels of selected genes in prostatic basal cells (n = 3) and prostatic luminal cells (n = 3). Boxes represent the interquartile range (IQR), center lines indicate the median, whiskers represent the data range, and individual data points are displayed as jittered dots. These plots illustrate differential expression patterns between the two prostate epithelial cell populations.

To evaluate the spatial expression and diagnostic utility of ITIH5 protein in clinical specimens, we performed immunohistochemical analysis on benign prostatic tissue, High-Grade Prostatic Intraepithelial Neoplasia (HGPIN), and prostatic adenocarcinoma, using p63 as a comparative standard. In histologically benign prostatic acini (Figure 2A), ITIH5 demonstrated robust staining that was strictly confined to the basal cell layer (Figure 2B). Importantly, this cytoplasmic ITIH5 staining is in the prostate basal cells that show nuclear basal cell staining of the established basal cell marker p63 (Figure 2C).

**Figure 2:**
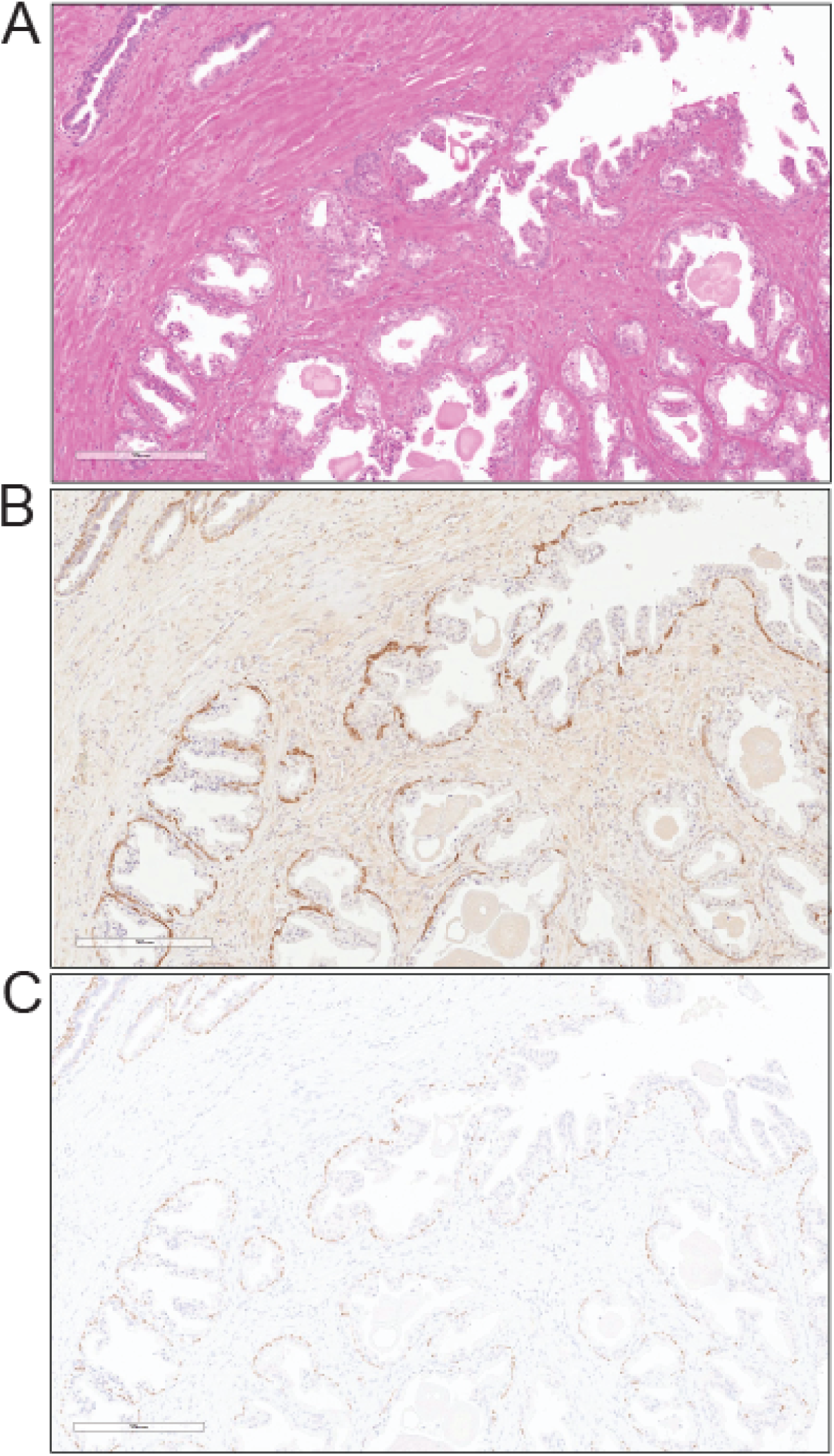
Immunohistochemical Localization of ITIH5 and p63 in Benign Prostatic Tissue. (A) Hematoxylin and eosin (H&E) staining of benign prostatic acini. (B) ITIH5 immunohistochemistry demonstrating strong, continuous staining restricted to the basal cell layer of the benign prostate. (C) p63 immunohistochemistry showing corresponding nuclear positivity in the basal cells of the same benign glands. These images confirm that ITIH5 specifically localizes to the basal cell compartment in morphologically benign prostate tissue.

We next examined ITIH5 expression in HGPIN, a recognized precursor to invasive carcinoma (Figure 3A). Both ITIH5 (Figure 3B) and p63 (Figure 3C) successfully highlighted the retained basal cell layer characteristic of HGPIN glands, distinguishing them from invasive disease.

**Figure 3:**
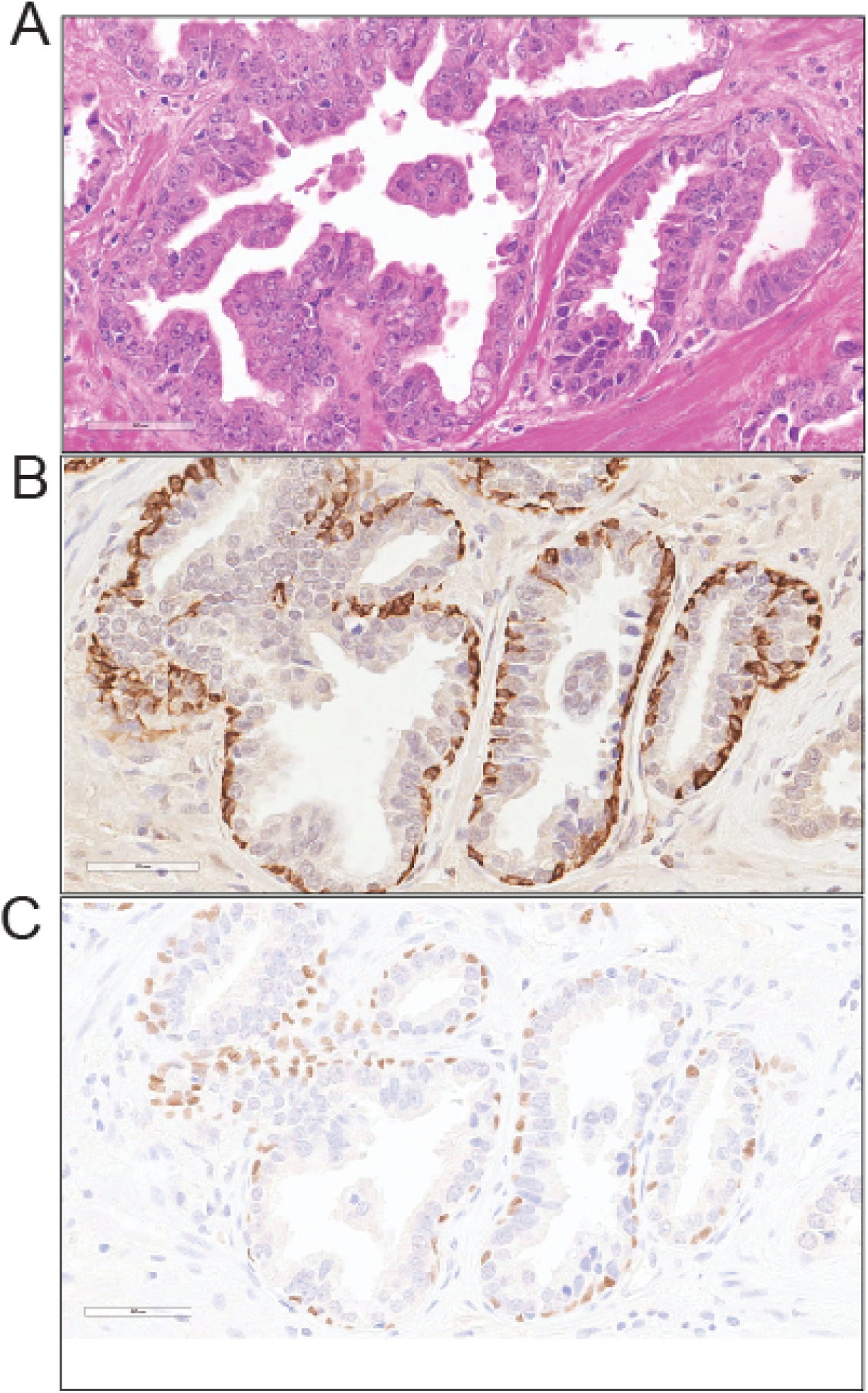
Expression of ITIH5 and p63 in High-Grade Prostatic Intraepithelial Neoplasia (HGPIN). (A) H&E staining showing glandular architecture and cytological atypia consistent with HGPIN. (B) ITIH5 immunostaining highlights the retained basal cells surrounding the atypical luminal proliferation in HGPIN. (C) p63 immunostaining confirms the presence of the basal cell layer in the same HGPIN glands. These images illustrate the utility of ITIH5 in identifying the intact basal cell layer in non-invasive precursor lesions.

Finally, to assess its behavior in malignancy, we evaluated areas where invasive prostate cancer directly juxtaposed benign glands (Figure 4A). The invasive adenocarcinoma glands exhibited a complete loss of ITIH5 expression, whereas the adjacent benign glands retained strong basal ITIH5 positivity (Figure 4B). This loss of ITIH5 in the malignant compartment was identical to the loss of p63 expression (Figure 4C). Together, these results confirm that ITIH5 is a highly specific basal cell marker whose absence characterizes invasive prostatic adenocarcinoma, suggesting it may serve as a valuable companion diagnostic marker alongside p63.

**Figure 4:**
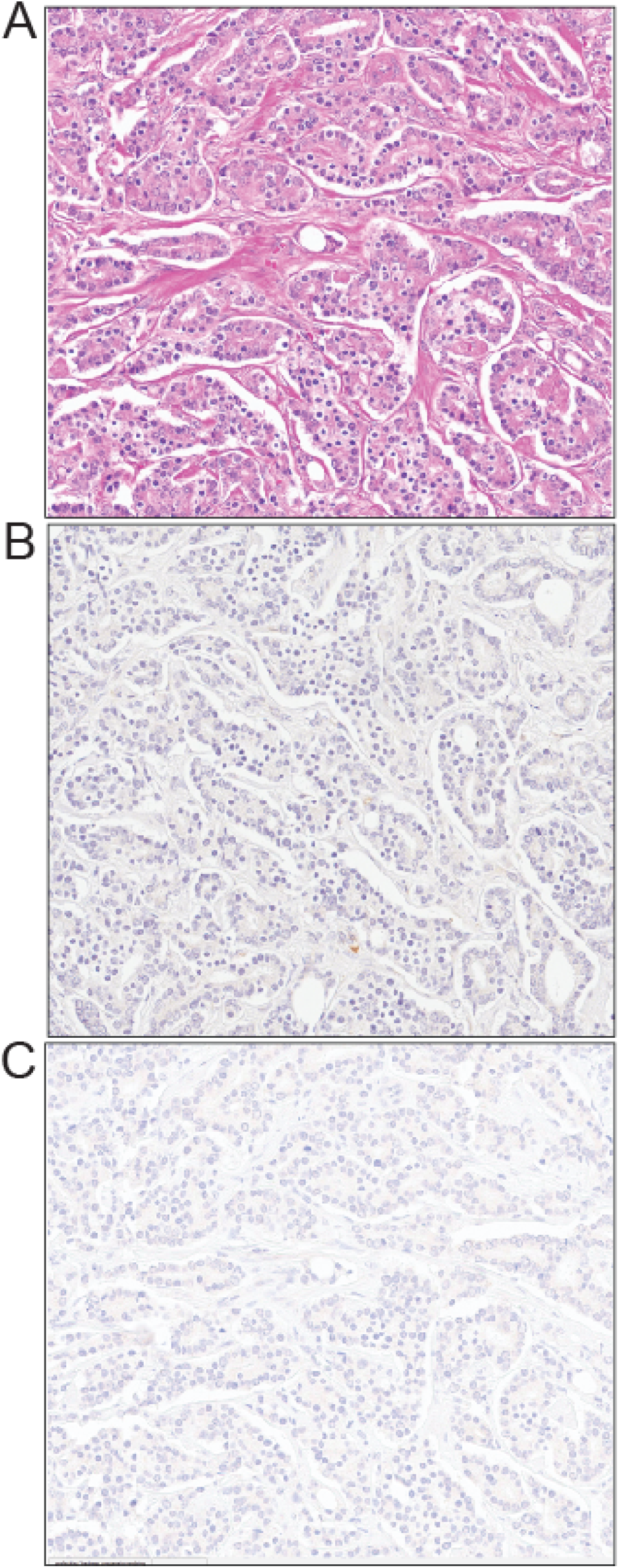
ITIH5 and p63 Expression in Invasive Prostatic Adenocarcinoma. (A) H&E staining depicting invasive prostate cancer glands infiltrating adjacent to a benign prostatic gland. (B) ITIH5 immunohistochemistry reveals a complete absence basal cell staining in the malignant glands, contrasting with the retained basal expression in the adjacent benign gland. (C) p63 staining correspondingly shows an absence due to absence of basal cells. This demonstrates the diagnostic utility of ITIH5 absence in invasive carcinoma cells, mirroring the established basal cell marker p63.

To further confirm the subcellular localization and specificity of ITIH5, we performed immunofluorescence staining on prostatic tissue sections (Figure 5). Consistent with the immunohistochemical findings, immunofluorescence revealed strong ITIH5 expression (green signal) strictly confined to the basal cell compartment. The signal outlines the periphery of the benign acini, while the inner secretory luminal cells and the surrounding stroma completely lack detectable ITIH5 expression. This orthogonal validation solidifies ITIH5 as a highly specific marker of the prostate basal cell layer.

**Figure 5:**
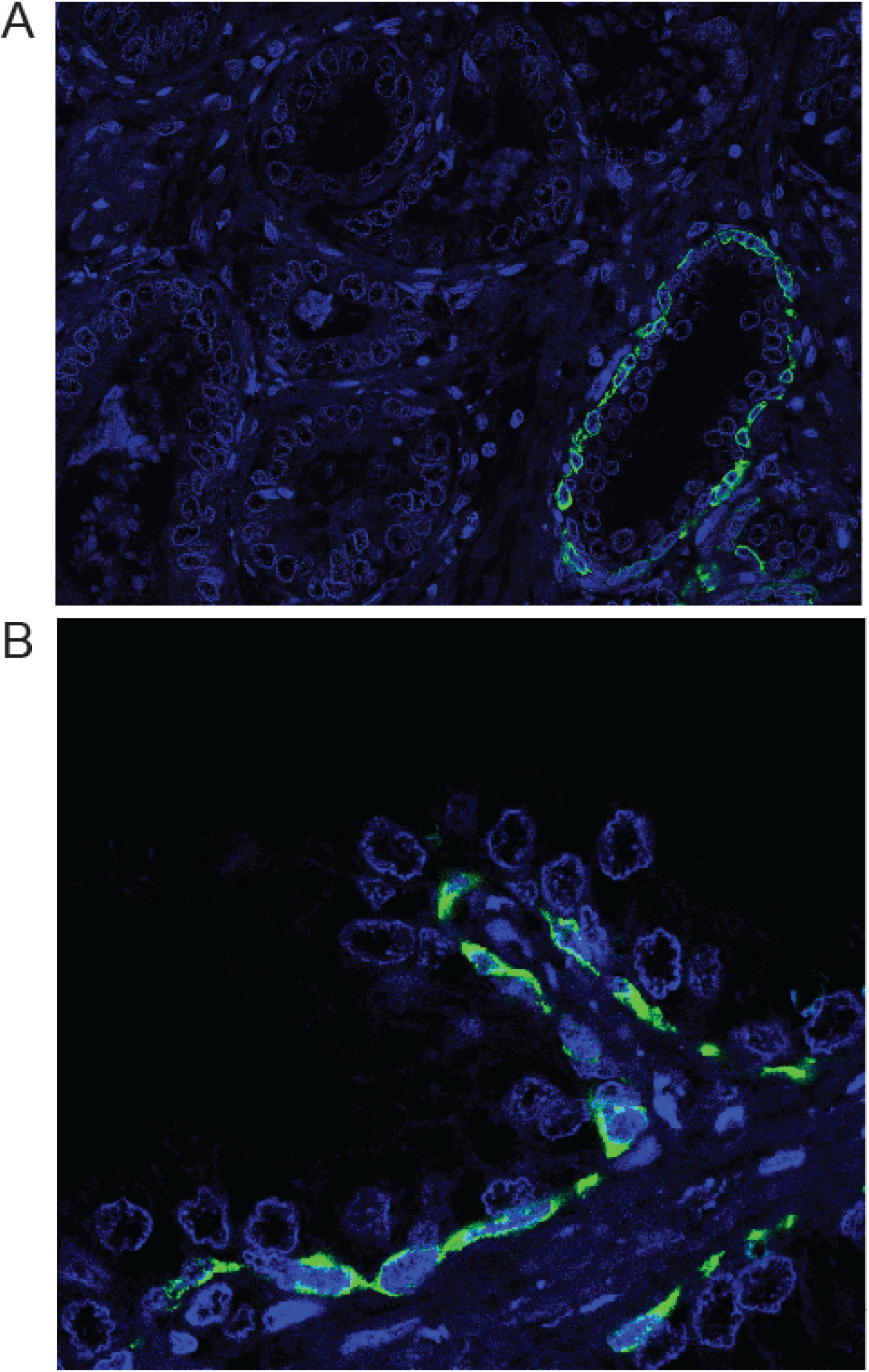
Immunofluorescence Localization of ITIH5 in Prostatic Acini. Representative immunofluorescence images of prostatic tissue stained for ITIH5 (green) and counterstained with DAPI for nuclei (blue). Both the top and bottom panels demonstrate robust, specific ITIH5 expression localized exclusively to the basal cell layer. The fluorescent signal is absent in the inner secretory luminal cells and the surrounding stroma.

## Discussion

The observation that ITIH5 strictly localizes to the basal cell compartment, without staining the adjacent secretory luminal cells, introduces a potentially valuable addition to the diagnostic immunohistochemical repertoire. Establishing the presence of a continuous or fragmented basal cell layer is the cornerstone of excluding invasive prostatic adenocarcinoma in histologically equivocal lesions (8–10). Currently, staining with p63 and 34βE12 cocktails is commonly employed to overcome the biological and technical variability that can result in the failure to detect basal cells when using a single marker (9). A novel marker such as ITIH5, which demonstrates a high fidelity for basal cells, could serve as a useful adjunct in cases where traditional marker expression is ambiguous or technically compromised. Furthermore, its robust and distinct cytoplasmic staining pattern makes ITIH5 an excellent candidate for integration into digital pathology workflows, where it could be evaluated via continuous scoring algorithms on open-source platforms like QuPath to automate and standardize tissue analysis. Future studies evaluating ITIH5 in common diagnostic mimickers, such as basal cell hyperplasia, partial atrophy, or seminal vesicle epithelium, will also be necessary to fully define its practical utility in routine sign-out.

The strict basal expression pattern of ITIH5 mirrors the utility of other emerging prostatic biomarkers. For instance, TRIM29 and ETS2 have recently been identified as being selectively expressed in benign prostate basal cells, with significantly down-regulated or absent expression in usual-type prostatic adenocarcinomas(5,10). Interestingly, while normal prostatic glands uniformly express basal markers, rare variants of prostate cancer can aberrantly express p63 and exhibit an intermediate immunophenotype between basal and luminal cells(8). Because a specific subset of tumors can sometimes unexpectedly retain p63 expression on immunohistochemistry, it will be critical for large-cohort studies to monitor whether any rare prostate carcinoma variants aberrantly retain ITIH5. The distinct luminal-sparing staining pattern of ITIH5 suggests it may play a specialized role in maintaining the basal epithelial phenotype. Consistent with this notion, ITIH5 re-expression in basal-type MDA-MB-231 breast cancer cells shifted the integrin profile towards β1-integrin, up-regulated the basement membrane collagens COL4A1 and COL4A2 as well as the desmosomal components desmocollin-2 and desmoplakin, and induced the formation of epithelial-like cell clusters (16). Moreover, a cytoplasmic ITIH5 staining pattern predominantly localized to the basal cell layer has previously been described in normal squamous epithelium of the uterine cervix, with decreased expression in high-grade precursor lesions and loss in invasive carcinoma (17), indicating that basal cell expression of ITIH5 may not be restricted to the prostate. Further investigation into its expression in atypical glandular proliferations and rare basal-like carcinomas will be critical to fully elucidate its diagnostic and biological significance.

The loss of basal markers in primary prostate cancer can be linked to early genetic events, such as TMPRSS2-ERG fusions. ERG overexpression alters 3D chromatin architecture, which decreases enhancer activity at distal regulatory elements and suppresses master basal regulators like Trp63 (25). If ITIH5 is similarly down-regulated by ERG-mediated chromatin remodeling, it would suggest that ITIH5 loss is an integral part of the epigenetic reprogramming that extinguishes the basal cell layer during early prostate carcinogenesis. Given its role in extracellular matrix stability(13,18), ITIH5 may also serve as a structural guardian at the basal-stromal barrier. Rather than acting as a protease inhibitor itself (within the inter-α-inhibitor complex, the Kunitz-type serine protease-inhibitory activity resides in the light chain bikunin, which is expressed almost exclusively in the liver (26), robust basal ITIH5 expression could support this barrier by stabilizing the pericellular hyaluronan matrix and modulating cell-matrix adhesion (11,13) (23,25). In prostate invasive carcinoma, the absence of ITIH5 most likely reflects the loss of the basal cell layer itself rather than epigenetic silencing within tumor cells, as luminal cells do not express ITIH5. Whether ITIH5 is additionally silenced by promoter hypermethylation in prostate cancer, as shown in breast, bladder, colon, lung and cervical cancer(11) (15–17,26), remains to be determined. Loss of the ITIH5-expressing basal cell layer may compromise this barrier and facilitate the basement membrane degradation required for stromal invasion by malignant luminal cells. Simultaneously, the tumor microenvironment undergoes profound remodeling during this invasive process. For example, the low-affinity nerve growth factor receptor (p75), which is normally restricted to benign basal cells, is progressively lost in the neoplastic epithelium but aberrantly re-expressed in stromal smooth muscle cells surrounding malignant foci (27). Evaluating the loss of epithelial ITIH5 alongside stromal indicators like p75 could provide a comprehensive assessment of the basal-stromal junction, aiding in complex differential diagnoses of atypical small acinar proliferations. Finally, while these findings establish a strong foundational role for ITIH5 in prostate pathology, this study is limited by the sample size of the clinical specimens evaluated. Broader validation across diverse histological grades, including higher Gleason patterns and specific morphological variants like ductal adenocarcinoma, is required to cement ITIH5’s diagnostic and biological significance.

## Acknowledgment

We would like to thank John Ness from the UAB pathology tissue biorepository for help with immunohistochemistry. This research supported by the UAB High Resolution Imaging Facility. We thank Sonja von Serenyi (Institute of Pathology, RWTH Aachen University) for her excellent technical assistance in the characterization of the ITIH5 antibody

## Data availability statement

The RNA-seq data used in this manuscript is from a publicly available data source from GEO.

